# Ageing attenuates exercise-enhanced motor cortical plasticity

**DOI:** 10.1101/2023.08.14.553314

**Authors:** Dylan Curtin, Claire J. Cadwallader, Eleanor M. Taylor, Sophie C. Andrews, Julie C. Stout, Joshua J. Hendrikse, Trevor T-J. Chong, James P. Coxon

## Abstract

Cardiorespiratory exercise is known to modulate motor cortical plasticity in young adults, but the influence of ageing on this relationship is unknown. Here, we compared the effects of a single session of cardiorespiratory exercise on motor cortical plasticity in young and older adults. We acquired measures of cortical excitatory and inhibitory activity of the primary motor cortex using transcranial magnetic stimulation (TMS) from 20 young (*M* ± s.d. = 25.30 ± 4.00 years) and 20 older (*M* ± s.d. = 64.10 ± 6.50 years) healthy adults. Single and paired pulse TMS measures were collected before and after a 20-minute bout of high-intensity interval cycling exercise or an equivalent period of rest, and again after intermittent theta burst stimulation (iTBS). In both young and older adults, exercise led to an increase in glutamatergic excitation and a reduction in gamma-aminobutyric acid (GABA) inhibition. However, in contrast to younger adults, older adults showed an attenuated plasticity response to iTBS following exercise. These results demonstrate an age-dependent decline in cortical plasticity and indicate that a preceding bout of high-intensity interval exercise may be less effective for enhancing primary motor cortex plasticity in older adults. Our findings align with the hypothesis that the capacity for cortical plasticity is altered in older age.

**Key points:**

- Exercise enhances motor cortical plasticity in young adults, but how ageing influences this effect is unknown.
- Here, we compared primary motor cortical plasticity responses in young and older adults before and after a bout of high-intensity interval exercise, and again after a plasticity-inducing protocol – intermittent theta burst stimulation.
- In both young and older adults, exercise led to an increase in glutamatergic excitation and a reduction in gamma-aminobutyric acid (GABAergic) inhibition.
- Our key result was that older adults showed an attenuated plasticity response to theta burst stimulation following exercise, relative to younger adults.
- Our findings demonstrate an age-dependent decline in exercise-enhanced cortical plasticity and indicate that a preceding bout of high-intensity interval exercise may be less effective for enhancing primary motor cortex plasticity in older adults.

## Introduction

The motor system is known to deteriorate with age (Seidler *et al*., 2010). The primary motor cortex is a key node within the motor system, which demonstrates plasticity as a function of use and learning (Rioult-Pedotti *et al*., 1998; Sanes & Donoghue, 2000; Levin *et al*., 2014). It is known to be a critical brain region for the acquisition and retention of motor skills (Muellbacher *et al*., 2002; Coxon *et al*., 2014; Peters *et al*., 2017; Kolasinski *et al*., 2019). Two major imperatives of motor ageing research have been to understand the neurophysiological mechanisms underlying age-related deterioration (Bhandari *et al*., 2016) and determine ways to mitigate or slow these declines (Semmler *et al*., 2021; Erickson *et al*., 2022).

The effects of ageing on primary motor cortex neurophysiology have predominantly been investigated using transcranial magnetic stimulation (TMS) (Bhandari *et al*., 2016; Semmler *et al*., 2021). Paired pulse TMS studies have demonstrated age-related alterations in inhibitory (Peinemann *et al*., 2001; Marneweck *et al*., 2011; Mooney *et al*., 2017; Hermans *et al*., 2018; Cuypers *et al*., 2020) and excitatory cortical networks (McGinley *et al*., 2010; Opie *et al*., 2020). Such changes are also known to limit the capacity for motor cortical plasticity in older age, as tested by various forms of repetitive TMS (Muller-Dahlhaus *et al*., 2008; Zimerman & Hummel, 2010; Freitas *et al*., 2011; Bhandari *et al*., 2016). Although these TMS studies indicate age-related declines in motor cortical networks, there has been limited investigation into how these changes may be mitigated or slowed (Semmler *et al*., 2021).

There is compelling evidence that cardiorespiratory exercise may be an effective way to modulate motor cortical plasticity. In healthy young adults, acute exercise, particularly when performed as high-intensity intervals, enhances motor cortical plasticity (Mang *et al*., 2014; Andrews *et al*., 2020) and modulates the balance between motor cortical excitation and inhibition, referred to as the E:I balance (Singh *et al*., 2014; Smith *et al*., 2014; Mooney *et al*., 2016; Neva *et al*., 2017; Stavrinos & Coxon, 2017; Coxon *et al*., 2018; Morris *et al*., 2020; Curtin *et al*., 2023). In older adults, there is strong evidence to support the efficacy of long-term exercise for attenuating age-related declines in motor function (Di Lorito *et al*., 2021; Valenzuela *et al*., 2023) and brain atrophy (Erickson *et al*., 2011; Maass *et al*., 2015; Ji *et al*., 2021), although the mechanisms underlying these long-term benefits are unclear. Using single and paired pulse TMS, a recent study (Neva *et al*., 2022) showed that acute exercise increased primary motor cortical excitation among a group of older adults, implicating the primary motor cortex and providing further support for the use of exercise to modulate the ageing motor system. Importantly, however, this study lacked a young comparison group and did not administer a plasticity-inducing protocol, which limits any conclusions about the effects of ageing on exercise-induced changes in motor cortical plasticity. This is important, as previous research has demonstrated reduced motor cortex plasticity in ageing (see e.g., Muller-Dahlhaus *et al*., 2008; Zimerman & Hummel, 2010; Freitas *et al*., 2011), which has the potential to attenuate the acute effects of exercise.

Here, we compared the effects of high-intensity interval exercise on excitatory and inhibitory cortical networks following intermittent theta burst stimulation (iTBS) in young (18–33 years) and older (51–74 years) adults. iTBS induces a long-term potentiation (LTP)-like effect through the application of brief, high-frequency subthreshold trains of stimulation in the gamma frequency band (50 Hz), superimposed upon a theta rhythm (5 Hz) (Huang *et al*., 2005; Suppa *et al*., 2016). Our group has previously shown that iTBS is sensitive to the effects of exercise on motor cortical networks (Andrews *et al*., 2020; Andrews *et al*., 2022). We predicted that relative to younger adults, older adults would show an attenuated plasticity response following exercise and iTBS, as quantified by smaller changes in the excitation:inhibition balance (i.e., smaller reductions in cortical inhibition and smaller increases in cortical facilitation).

### Participants

## Methods

We tested 20 younger (*M* ± s.d. = 25.30 ± 4.00 years) and 20 older adults (*M* ± s.d. = 64.10 ± 6.50 years; Table 1). Our sample included data from 12/20 of the young adults and 4/20 older adults reported in our previous study (Andrews *et al*., 2020). Exclusion criteria included age < 18 or > 35 years for younger adults and < 50 or > 75 years for older adults; personal history of neurological disease or moderate-severe psychiatric illness; chronic or recent use of drugs of addiction; age-related cognitive impairment (Montreal Cognitive Assessment < 26) (Nasreddine *et al*., 2005); and any contraindications to TMS (Rossi *et al*., 2009) or high-intensity exercise (Sports Medicine Australia, 2011). Three older participants were taking a low dose of an antidepressant (selective serotonin reuptake inhibitor), otherwise, participants were free of psychotropic medications. Groups did not differ on key demographic and health measures (Table 1). All participants provided written informed consent. The experimental procedures were conducted in accordance with the most recent version of the Declaration of Helsinki, except for registration in a database, and were approved by the Monash University Human Research Ethics Committee (Project reference #16567).

**Table 1.** Participant characteristics.

| Characteristics | Younger | Older | Test statistic |
| --- | --- | --- | --- |
| *n* | 20 | 20 | - |
| Age, years (range) | 25.30 ± 4.00 (18–33) | 64.10 ± 6.50 (51–74) | $t(38) = 22.74, p < .0001$ |
| Sex, F/M | 14F / 6M | 11F / 9M | $\chi^2(1) = 0.96, p = .327$ |
| Body mass index, range | 22.94 ± 3.18 | 26.15 ± 3.63 | $t(38) = 2.98, p = .005$ |
| Formal Education, years | 16.50 ± 2.73 | 16.33 ± 3.52 | $t(36) = 0.16, p = .871$ |
| Handedness, EHI | 81.88 ± 19.39 /<br>1 Left -36.36 | 81.84 ± 15.84 | $t(36) = 0.70, p = .491$ |
| Physical activity levels,<br>IPAQ, MET minutes/week | 5024 ± 4957 | 5683 ± 389 | $t(36) = 0.45, p = .654$ |
| Hospital Anxiety<br>Depression Scale (HADS) | 8.50 ± 5.73 | 6.35 ± 4.34 | $t(35) = 0.21, p = 0.214$ |
| Blood pressure,<br>Systolic/Diastolic | - | 133.19 ± 12.76 /<br>81.64 ± 7.37 | - |
| Montreal Cognitive<br>Assessment, MoCA | - | 27.94 ± 1.00 | - |
| VO<sub>2</sub> Peak (mL.kg<sup>-1</sup>.min<sup>-1</sup>)<sup>a</sup> | - | 38.22 ± 12.29 | - |

*Note.* Values are means ± standard deviation. *n* = number of participants, F = female, M = male, EHI = Edinburgh handedness inventory [range: -100 (strongly left-handed), 0 (ambidextrous), +100 (strongly right-handed)] (Oldfield, 1971), IPAQ = international physical activity questionnaire (higher scores indicate higher physical activity levels) (Craig *et al.*, 2003), MET = metabolic equivalent, HADS scores represent the summed scores of depression and anxiety subscale, VO<sub>2</sub> peak = highest amount of oxygen consumed at peak exercise.
<sup>a</sup> = Data obtained from 16 older adults

### Experimental Design

Participants completed two sessions involving TMS and either exercise or rest. Due to the tendency for many age-based equations to underestimate maximum heart rate in the elderly (Nes *et al*., 2013), older adults (*n* = 16/20) also completed an incremental cardiorespiratory exercise assessment, prior to the TMS sessions, so that exercise could be more accurately prescribed in TMS sessions (Figure 1A). All sessions were separated by a washout period of >72 hours to limit any carry-over effects of exercise or non-invasive brain stimulation. The order of sessions two and three was counterbalanced according to a pre-determined pseudo-randomisation scheme. Participants were instructed to refrain from moderate and high intensity exercise in the 24 hours before any session.

**Figure 1.**
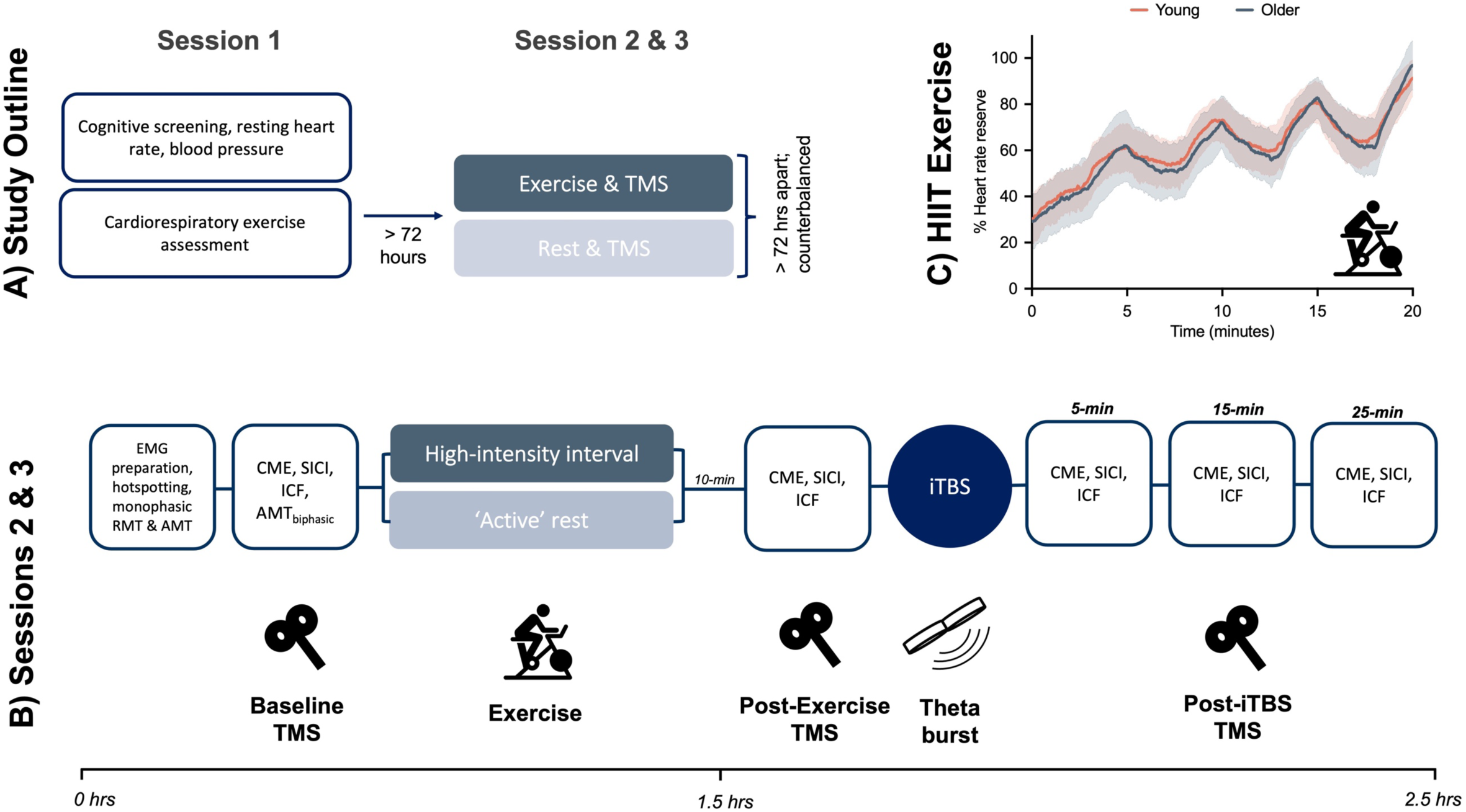
Panel. **A)** Older adults completed an initial session involving cognitive screening and a cardiorespiratory exercise assessment, i.e., Session 1. All participants completed two transcranial magnetic stimulation (TMS) sessions, one involving exercise and the other rest, i.e., Sessions 2 and 3. **Panel B)** In sessions 2 and 3, participants underwent baseline TMS measures of excitatory and inhibitory cortical activity, followed by a 20-minute bout of high-intensity interval exercise or rest. Repeat TMS measures were taken after a 10-minute break, and then again 5, 15, and 25 minutes after intermittent theta burst stimulation (iTBS). **Panel C)** Percent heart rate reserve during high-intensity interval exercise (HIIT) in young (orange) and older (dark blue) adults, with group data presented as mean ± 1 standard deviation. *Note*. EMG = electromyography, RMT = resting motor threshold, AMT = active motor threshold, CME = cortico-motor excitability, SICI = short-interval intracortical inhibition, ICF = intracortical facilitation.

Figure 1B shows an overview of the sequence of events for TMS sessions. Baseline TMS measures of excitatory and inhibitory cortical activity were taken. Participants then completed 20 minutes of high-intensity interval cycling or active rest on a stationary ergometer, followed by a 10-minute cool-down period. Repeat TMS measures were taken before and after iTBS at 5-, 15-, and 25-minute intervals. TMS sessions were scheduled at the same time of day to reduce intraindividual diurnal effects on TMS responses (Ridding & Ziemann, 2010) or exercise performance.

### Cardiorespiratory exercise assessment

Older adults completed an incremental cardiorespiratory exercise assessment on a stationary cycle ergometer (Wattbike Trainer, Geelong, Australia), integrated with a cardio-pulmonary monitoring system (ADInstruments, Dunedin, New Zealand). All tests were overseen by an experienced exercise scientist (J.P.C.). Before beginning the protocol, blood pressure (Omron HEM-7320, Omron, Kyoto, Japan) and resting heart rate (Polar H10 heart rate monitor) were assessed while participants were seated comfortably. Maximum heart rate was estimated as 220 – (0.7 x Age in years) (Tanaka *et al*., 2001). From this, we estimated heart rate reserve, calculated as the difference between maximum heart rate and resting heart rate.

After a brief warm-up, participants cycled for 3 minutes at 40% heart rate reserve followed by 3 minutes at 60% heart rate reserve. Thereafter, participants were instructed to maintain a steady cadence while power was increased by 10–15 watts each minute. Participants were verbally encouraged to carry on for as long as they could, and the test was terminated when the participant reached volitional exhaustion, with a goal of 8–12 minutes of cycling time in total. At the end of each minute, participants were asked to rate their level of perceived exertion, using Borg’s perceived exertion scale, a self-report measure ranging from 6 (no exertion at all) to 20 (maximal exertion) (Borg, 1970).

The following exercise response measures were obtained and analysed offline: oxygen uptake (litre per minute), carbon dioxide uptake (litre per minute), maximal oxygen volume uptake recorded at peak exercise (VO_2_ peak, in mL.kg^-1^.min^-1^), respiratory exchange ratio (volume of VO_2_ consumed/VCO_2_ produced), and heart rate. For a maximal effort to be verified, two of three criteria needed to be met, including: 1) a respiratory exchange ratio ≥ 1.05 for people aged 50–64 years, or ≥ 1.00 for people aged 65 years and above; 2) a heart rate > 95% of heart rate reserve; 3) self-rated exertion level ≥ 17 (Borg scale) (Edvardsen *et al*., 2014). Seven participants did not meet these criteria. For these participants, individual regression equations were used (mean *R*^2^ = .96) to derive an estimated VO_2_ peak, based on their sub-maximal values (Hendrikse *et al*., 2021).

From the cardiorespiratory exercise assessment, we calculated each participant’s VO_2_ reserve, defined as the difference between resting VO_2_ and VO_2_ peak, and the relationship between VO_2_ reserve and HR reserve (Pearson’s *r* = .99, *p* < .0001). Based on this relationship, exercise HR targets were calculated for subsequent exercise (e.g., heart rate at 90 % VO_2_ reserve).

### High-intensity interval exercise protocol

High-intensity interval exercise was performed on a stationary cycle ergometer (Wattbike Atom, Geelong, Australia). Our protocol consisted of alternating intervals of 3-min of cycling at 50% heart rate reserve, and 2-min of cycling at up to ∼90% heart rate reserve, for a total duration of 20 minutes (Figure 1C) (Stavrinos & Coxon, 2017; Andrews *et al*., 2020). For younger participants, we calculated an estimated heart rate reserve using resting heart rate assessed during the first session whilst seated comfortably and estimated maximum heart rate using the formula: 220 – (0.7 x Age in years) (Tanaka *et al*., 2001).

Throughout exercise, we quantified participant ratings of exercise intensity using Borg’s perceived exertion scale (Borg, 1970). We instructed participants to refrain from gripping the bike handlebar during exercise (Andrews *et al*., 2020; Curtin *et al*., 2023). Participants completed a 2-minute warm-up and 2-minute cool-down period by cycling at a very low intensity. After completing exercise, participants rested for 10 minutes.

### Rest Protocol

We chose a control rest protocol that was designed to account for posture and lower limb movement across sessions. In this protocol, participants sat on the stationary exercise bike and were instructed to turn the pedals over at a very low cadence for 20 minutes (Cadwallader *et al*., 2023). The goal of this session was to keep the participant’s heart rate within the range of 10 beats per minute from their resting heart rate [(achieved mean difference for younger adults = 10.82, older adults = 6.76; independent samples *t* test, *t*(20) = 1.13, *p* = .270].

### Transcranial magnetic stimulation

For TMS, participants were seated with both hands resting in a pronated position on a pillow in their laps. We recorded surface electromyography (EMG) from the first dorsal interosseous muscle of the dominant hand using Ag-AgCl electrodes (Ambu White Sensor WS, Ambu, Copenhagen, Denmark), arranged in a belly-tendon montage and secured with sports tape for the duration of the session. Raw EMG signals were amplified 1000x, bandpass filtered at 10– 1000 Hz (Digitimer D360 amplifier, Digitimer Ltd, Welwyn Garden City, England), digitised at 2000 Hz, and stored for offline analysis using Signal (Version 6.05a). Each EMG sweep began 100 ms before the TMS trigger and finished 200 ms after the trigger.

Monophasic TMS pulses were applied to the dominant primary motor cortex using a figure-of-eight coil (70 mm wing diameter), connected to two Magstim 200^2^ magnetic stimulators via a Bistim^2^ module (Magstim Company Ltd, Whitland, Wales). We identified the optimal site to elicit a consistent motor evoked potential (MEP) in the dominant FDI (the ‘hot-spot’) and marked this spot to ensure a consistent coil position across each session. The TMS coil was positioned tangentially to the skull with the handle orientated posteriorly, at approximately 45° to the central sulcus, resulting in a posterior-anterior current flow in the cortex. Resting motor threshold (RMT) was defined as the lowest stimulation intensity required to elicit motor evoked potentials (MEPs) of ≥ 50 μV in the relaxed FDI in at least 5 out of 10 consecutive trials. Active motor threshold (AMT) was defined as the lowest stimulation intensity required to elicit MEPs of ≥ 200 μV during a weak tonic contraction of the FDI in at least 5 out of 10 consecutive trials (Rossi *et al*., 2009).

To examine primary motor cortex neurophysiology, we used both single-pulse TMS (corticomotor excitability, CME), and paired-pulse TMS (short-interval intracortical inhibition, SICI; and intracortical facilitation, ICF). The intensity of a single test stimulus was adjusted to produce a stable MEP of approximately 1mV at baseline. For paired-pulse TMS, we used a sub-threshold conditioning stimulus that preceded this supra-threshold test stimulus by 2 ms and 12 ms, respectively. For SICI, the conditioning stimulus intensity was adjusted to produce 50 % inhibition of the non-conditioned MEP at baseline. The same conditioning and test stimulus intensities were used for ICF. To ensure valid comparison of paired-pulse measures at the post-exercise time point, the test stimulus intensity was adjusted, where necessary, so that test MEP sizes were matched to within 30 % of the Baseline time-point (Andrews et al., 2020). Single- and paired-pulse measures were delivered in a pseudorandomised order at 6 second intervals (± 15 % jitter) at each time point.

### Intermittent theta burst stimulation (iTBS)

iTBS was applied to the primary motor cortex, using a C-B60 figure-of-eight coil (70 mm diameter) connected to a MagVenture MagPro X100 stimulator (MagVenture Ltd.). The iTBS protocol followed standard procedures, i.e., three 50 Hz biphasic pulses delivered every 200 ms for 2 s, repeated every 10 s for 20 repetitions (Huang *et al*., 2005). Stimulation intensity was set at 80% of biphasic AMT. To minimise confounding effects of prior voluntary FDI muscle activity on iTBS (Huang *et al*., 2007), we measured biphasic AMT at the end of the baseline time-point (Figure 1B).

### TMS analysis

Exercise has the potential to influence MEP amplitudes through an effect on background muscle activity. We addressed this potential confound by inspecting EMG recordings offline and discarded trials contaminated with muscle activity in the 100 ms before the TMS pulse, or with a root-mean-square value ≥ 10 μV over this period. This resulted in the removal of 2.37% and 6.71% of trials in the young and older adults in the rest session, respectively, and 1.38% and 6.94% of trials in the HIIT session. For all single- and paired-pulse trials, the peak-to-peak amplitude of each MEP was obtained and averaged at each time point. For paired-pulse measures, the amplitude of the conditioned MEP was expressed as a ratio of the amplitude of the non-conditioned MEP (C/NC). To reduce the influence of outliers on central tendency, we used an *a priori* trimmed mean procedure, in which the highest and lowest value of each set of 16 MEPs was discarded (Coxon *et al*., 2014; Andrews *et al*., 2020; Curtin *et al*., 2023).

To characterise the combined effects of excitatory and inhibitory cortical activity, we calculated an excitation:inhibition (E:I) ratio (Andrews *et al*., 2020; Curtin *et al*., 2023) using data obtained from SICI and ICF measures. An increase in the E:I value represents a combination of increased excitation and reduced inhibition.

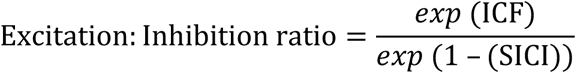

To assess the effects of age on cortical responses to exercise and theta burst stimulation, we used mixed analysis of variance (ANOVA), with Age as a between-subjects factor and Exercise Session and Time as within-subject factors. The values of 16 univariate outliers (0.0008 % of all valid TMS trials), assessed using a criterion of *z* = ± 3.29, were adjusted to one unit higher than the next most extreme value in their respective conditions (Tabachnick & Fidell, 2013). Mixed model ANOVAs were conducted for each of our measures (CME, SICI, ICF, E:I balance), and significant effects were followed-up using Bonferroni-adjusted pairwise comparisons. To assess the magnitude of effects, partial eta-squared (17_p_^2^) and Cohen’s *d* were calculated, where appropriate. To account for differences in some baseline TMS values between age groups (see Table 3), we used delta change scores for age comparisons, calculated as the difference between the post-exercise or post-iTBS timepoint and baseline. Data were analysed using JASP version 0.17.1 with alpha set at .05.

### Exercise analysis

During high-intensity exercise and active control sessions, heart rate, power (Watts), and cadence were collected at 1-second intervals. Ratings of perceived exertion were collected at the end of each minute of exercise during HIIT, and every 5 minutes during the active control condition. Heart rate data from 5 older adult sessions (two in the Rest session, and three in the HIIT session) were omitted due to technical difficulties with our heart rate monitor.

### Data availability

The data reported in this study are openly available via FigShare at http://doi.org/10.26180/23622423

## Results

Two older adults with pre-existing muscle or joint pain experienced mild discomfort related to their conditions during exercise. All effects were transient and did not necessitate stopping exercise. No other adverse outcomes related to exercise, or TMS, were reported.

### Exercise performance

Participants exercised at the intended heart rate intensity (Figure 1C; Table 2), indicating that our exercise intervention achieved the desired physiological stimulus. There were no differences in heart rate intensity (% heart rate reserve), power:weight (Watts/kg), cadence, or perceived exertion between younger and older adults during either the exercise or active control condition (all *p* > .133).

**Table 2.** Exercise performance in young and older Adults.

| Session | %HRR |  | Power:Weight (Watts/kg) |  | Cadence (RPM) |  | RPE |  |
| --- | --- | --- | --- | --- | --- | --- | --- | --- |
| HIIT | Young | Older | Young | Older | Young | Older | Young | Older |
| 0-3 min | 44.05 ± 9.17 | 42.48 ± 12.28 | 0.98 ± 0.22 | 0.99 ± 0.32 | 70.51 ± 10.31 | 71.62 ± 10.92 | 10.00 ± 1.49 | 9.41 ± 1.37 |
| 4-5 min | 59.78 ± 9.54 | 60.29 ± 14.79 | 1.41 ± 0.33 | 1.55 ± 0.61 | 82.64 ± 11 | 82.10 ± 12.12 | 12.30 ± 1.13 | 11.82 ± 1.47 |
| 6-8 min | 54.25 ± 9.86 | 51.87 ± 11.41 | 0.98 ± 0.21 | 1.05 ± 0.29 | 69.71 ± 10.66 | 73.96 ± 10.29 | 11.10 ± 1.17 | 11.47 ± 1.62 |
| 9-10 min | 71.71 ± 8.75 | 69.11 ± 11.65 | 1.58 ± 0.42 | 1.75 ± 0.61 | 86.86 ± 11.78 | 85.35 ± 10.88 | 13.15 ± 1.39 | 13.18 ± 1.51 |
| 11-13 min | 60.04 ± 10.19 | 58.17 ± 10.47 | 0.99 ± 0.23 | 1.07 ± 0.31 | 70.97 ± 10.83 | 74.03 ± 10.52 | 11.40 ± 1.35 | 12.06 ± 1.52 |
| 14-15 min | 78.82 ± 8.55 | 79.31 ± 9.09 | 1.78 ± 0.56 | 1.94 ± 0.60 | 88.91 ± 12.67 | 88.34 ± 11.67 | 14.65 ± 1.39 | 14.88 ± 1.45 |
| 16-18 min | 64.09 ± 10.50 | 62.46 ± 11.62 | 0.99 ± 0.24 | 1.04 ± 0.32 | 70.47 ± 11.52 | 74.21 ± 9.97 | 12.15 ± 1.57 | 12.65 ± 2.00 |
| 19-20 min | 86.21 ± 9.30 | 91.19 ± 10.66 | 2.18 ± 0.79 | 2.55 ± 0.93 | 98.19 ± 12.75 | 96.92 ± 15.20 | 16.75 ± 1.41 | 17.53 ± 1.87 |
| **Control <sup>a</sup>** |  |  |  |  |  |  |  |  |
| 0-20 min | 8.88 ± 4.02 | 6.79 ± 9.82 | 0.04 ± 0.02 | 0.06 ± 0.03 | 17.21 ± 5.83 | 19.09 ± 5.36 | 7.66 ± 1.20 | 6.89 ± 1.10 |

Values are means ± standard deviation. HIIT = high-intensity interval training, RPM = revolutions per minute, RPE = rating of perceived exertion (Borg 6–20 Scale). For the HIIT session, %HRR, Power:Weight, and cadence were averaged across the last minute of each interval. Independent samples *t*-tests revealed no differences between groups on any measures of exercise performance in either the rest or HIIT session (all *p* > .133). <sup>a</sup> Control session includes data from 8 younger and 16 older adults.

**Table 3.** Baseline transcranial magnetic stimulation (TMS) characteristics by age category.

| TMS measure | Young ($n=20$) | Older ($n=20$) | Independent t-test |
| --- | --- | --- | --- |
| % TS/RMT | 120.47 ± 5.49 | 124.66 ± 6.18 | $t(38)$, $p = .029$ |
| % CS/AMT | 78.13 ± 6.67 | 81.73 ± 10.46 | $t(38)$, $p = .224$ |
| Biphasic AMT, %MSO | 39.39 ± 5.25 | 42.97 ± 8.28 | $t(38)$, $p = .111$ |
| iTBS, %MSO | 31.88 ± 4.04 | 34.28 ± 6.24 | $t(38)$, $p = .157$ |
| E:I balance | 1.86 ± 0.49 | 2.37 ± 0.95 | $t(38)$, $p = .037$ |
| SICI<sup>a</sup> | 0.48 ± 0.05 | 0.49 ± 0.07 | $t(38)$, $p = .611$ |
| ICF<sup>b</sup> | 1.08 ± 0.26 | 1.31 ± 0.40 | $t(38)$, $p = .036$ |
| CME | 1.03 ± 0.22 | 1.20 ± 0.54 | $t(38)$, $p = .204$ |

RMT = resting motor threshold, MSO = maximum stimulator output, AMT = active motor threshold, TS = test stimulus, CS = conditioning stimulus, iTBS = intermittent theta burst stimulation. Values are means ± standard deviation. As there were no main or interaction effects related to Session (all $p > .108$ ), data is averaged across the two sessions.
<sup>a</sup> = Separate rmANOVAs [2 Session x 2 TMS pulse (conditioned, non-conditioned)] for the two age groups confirmed the presence of inhibition at baseline, as indicated by a main effect of TMS pulse (Young and Older, both $p < .0001$ ).
<sup>b</sup> = Visual inspection of the ICF data indicated the presence of facilitation (i.e., an increased conditioned MEP amplitude relative to the NC response) in both age groups; however, the main effect of TMS pulse was not significant (Young, $p = .539$ , Older $p = .544$ ).

### Baseline transcranial magnetic stimulation (TMS)

There were no differences between younger and older participants on most baseline TMS measures (*p* > .111) (see Table 3). Older adults did show higher ICF (*p* = .036) and E:I values (*p* = .037) at baseline and higher 1mV test stimulus intensity relative to RMT compared to younger adults (*p* = .029).

### Excitation:inhibition balance (Figure 2A–C)

Our key result (Figure 2A) was a Session x Time x Age interaction for the delta E:I balance (*p* = .019; see Table 4 for full results). Bonferroni-adjusted pairwise comparisons showed that, compared with younger adults, older adults demonstrated a smaller increase in the E:I balance following iTBS in the HIIT session (*p* = .011, Cohen’s *d* = 0.85). There were no differences among age groups in the delta E:I balance following iTBS in the rest session (*p* = .680, Cohen’s *d* = 0.13), nor were there any differences between groups immediately following exercise (*p* = .532, Cohen’s *d* = 0.20) or rest (*p* = .715, Cohen’s *d* = 0.12).

**Figure 2.**
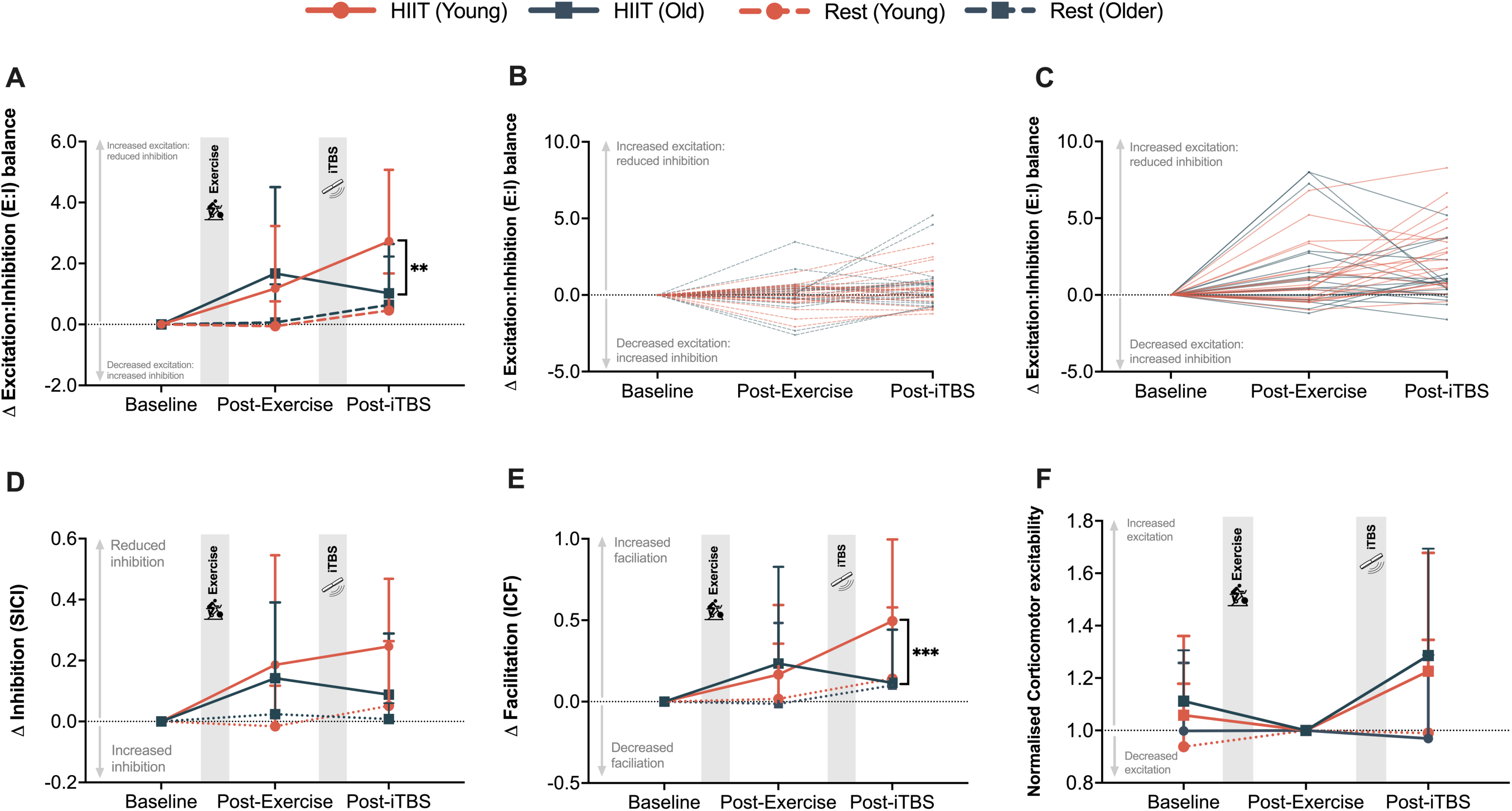
Age-related effects of exercise and intermittent theta burst stimulation on motor cortex excitation and inhibition. **(A)** Excitation:inhibition (E:I) ratio, which represents a combination of data obtained from SICI and ICF, with individual data plotted during the **(B)** active rest control and **(C)** HIIT session, **(D)** short-interval intracortical inhibition (SICI), (**E)** intracortical facilitation (ICF), and **(F)** corticomotor excitability. For panels A–E, the data is expressed as a delta change from baseline to Post-Exercise and the average of the three post-iTBS timepoints. For panel F, data is normalised to the Post-Exercise timepoint. Bolded lines represent the mean (+ 1 standard deviation). ** HIIT Younger Post-iTBS > HIIT Older Post-iTBS, *p* = .011, *** HIIT Younger Post-iTBS > HIIT Older Post-iTBS*, p* = .007

**Table 4.** Transcranial magnetic stimulation ANOVA results.

| Measure | Age | Session | Session x Age | Time | Time x Age | Session x Time | Session x Time x Age |
| --- | --- | --- | --- | --- | --- | --- | --- |
| E:I balance | $F(1,38) = 0.44, p$ | **$F(1,38) = 16.11, p$** | $F(1,38) = 1.24, p = .273,$ | **$F(1,38) = 5.47, p$** | **$F(1,38) = 6.31, p = .016$** | $F(1,38) = 0.04, p = .835$ | **$F(1,38) = 6.05, p = .019$** |
| | $= .513, \eta_p^2 = 0.01$ | **$< .0001, \eta_p^2 = 0.30$** | $\eta_p^2 = 0.03$ | **$= .025, \eta_p^2 = 0.13$** | **$p = 0.14$** | $\eta_p^2 = 0.001$ | **$\eta_p^2 = 0.14$** |
| SICI (C/NC) | $F(1,38) = 1.22, p$ | **$F(1,38) = 23.96, p$** | $F(1,38) = 2.64, p = .112,$ | $F(1,38) = 0.30, p$ | $F(1,38) = 3.43, p = .072,$ | $F(1,38) = 0.16, p = .694,$ | $F(1,38) = 0.08, p = .783,$ |
| | $= .276, \eta_p^2 = 0.03$ | **$< .0001, \eta_p^2 = 0.39$** | $\eta_p^2 = 0.07$ | $= .589, \eta_p^2 = 0.008$ | $\eta_p^2 = 0.08$ | $\eta_p^2 = 0.004$ | $\eta_p^2 = 0.002$ |
| ICF (C/NC) | $F(1,38) = 1.25, p$ | **$F(1,38) = 4.75, p$** | $F(1,38) = 0.47, p = .499,$ | **$F(1,38) = 6.77, p$** | **$F(1,38) = 7.05, p$** | $F(1,38) = 0.02, p = .885,$ | **$F(1,38) = 4.64, p = .038,$** |
| | $= .272, \eta_p^2 = 0.03$ | **$= .036, \eta_p^2 = 0.11$** | $\eta_p^2 = 0.01$ | **$= .013, \eta_p^2 = 0.15$** | **$= .012, \eta_p^2 = 0.15$** | $\eta_p^2 = 0.001$ | **$\eta_p^2 = 0.11$** |

Separate mixed model ANOVAs were performed on TMS measures using within-subject factors of Session (Rest, HIIT) and Time ( $\Delta$ Post-Exercise, $\Delta$ Post-iTBS) and a between-subjects factor of Age (Younger vs Older). For each time point, a delta value was calculated by subtracting the baseline time point (i.e., before exercise) from either Post-Exercise or Post-iTBS timepoints. E:I = excitation:inhibition balance, ICF = intracortical facilitation, SICI = short-interval intracortical inhibition. SICI and ICF data are expressed as a ratio of the amplitude of the conditioned (C) motor evoked potential to the amplitude of the non-conditioned (NC) motor evoked potential. Bolded values denote statistically significant results.

For the older adult group, a two-way rmANOVA revealed main effects of Session, *F*(1,19) = 6.13, *p* = .023, 17_p_^2^ = 0.24, and Time, *F*(2,38) = 6.04, *p* = .005, 17_p_^2^ = 0.24, but no Session x Time interaction, *F*(2,38) = 2.81, *p* = .073, 17_p_^2^ = 0.13. Pairwise comparisons showed that there was an increase in the E:I balance pre-to post-exercise (*p* = .006, Cohen’s *d* = 0.85). The E:I balance was also higher at the post-exercise time point in the HIIT compared to the rest session (*p* = .021, Cohen’s *d* = 0.92).

### Intracortical inhibition (Figure 2D)

We observed a main effect of Session for SICI (*p* < .001; Table 4), such that there was a greater release of inhibition in the HIIT compared to the rest session across both age groups (*p* < .001, Cohen’s *d* = 0.70). There were no other main or interaction effects (all *p* > .072; Table 4).

### Intracortical facilitation (Figure 2E)

Consistent with the E:I balance results, we observed a Session x Time x Age interaction for delta ICF (*p* = .038; Table 4). Bonferroni-adjusted pairwise comparisons showed that older adults showed a smaller increase in ICF following iTBS in the HIIT session compared to younger adults (*p* = .007, Cohen’s *d* = 0.90). The pattern of results is presented in Figure 3, showing representative MEPs for ICF from both a young and older participant. There were no differences among age groups in ICF following iTBS in the rest session (*p* = .737, Cohen’s *d* = 0.11), nor were there any differences between groups immediately following exercise (*p* = .681, Cohen’s *d* = 0.13) or rest (*p* = .819, Cohen’s *d* = 0.07).

**Figure 3.**
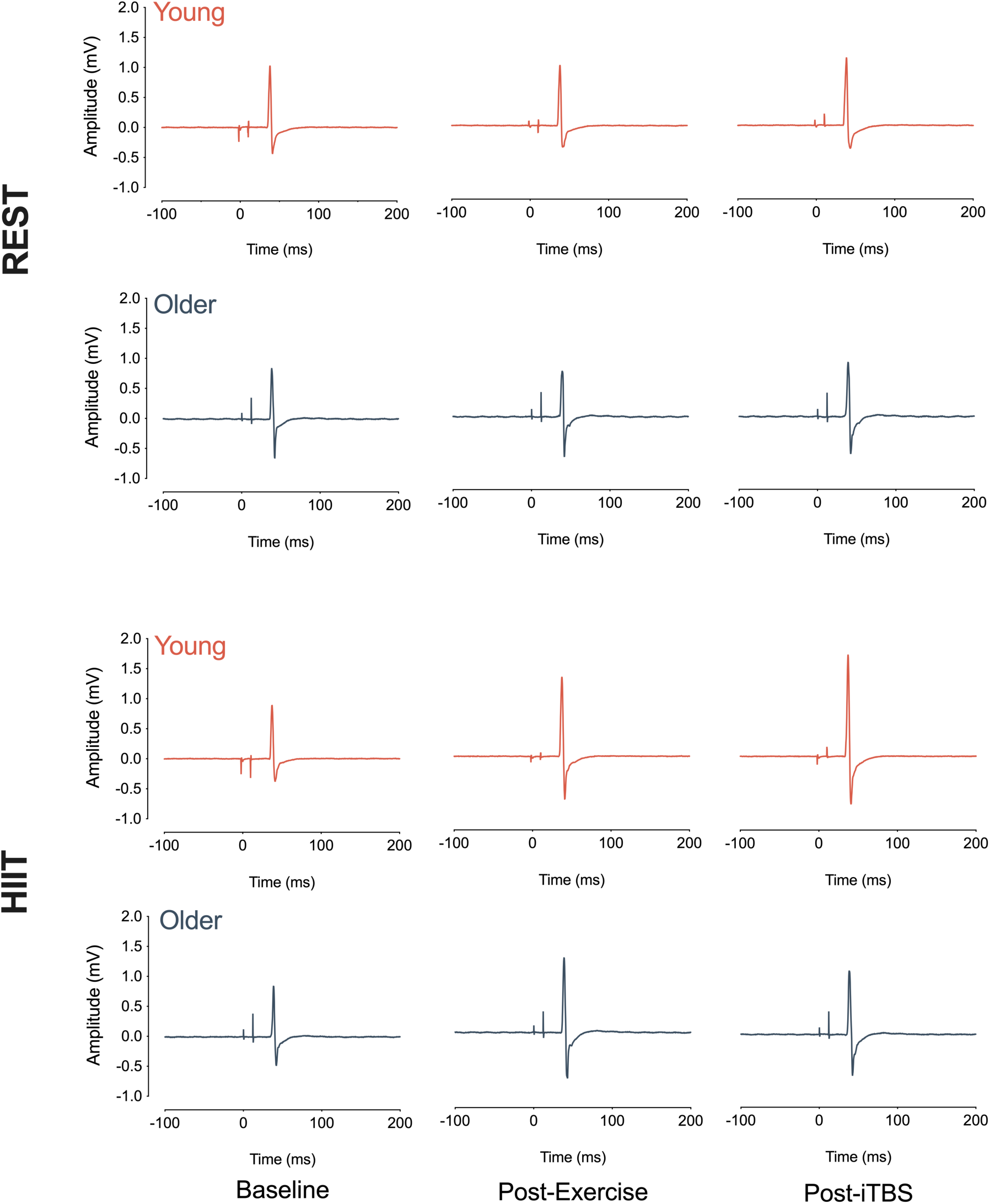
Motor evoked potentials (MEPs) for paired-pulse TMS with 12 ms inter-stimulus interval (ICF). Examples are from a representative Younger (orange) and Older (dark blue) participant within the rest (top panels) and high-intensity interval (HIIT) exercise (bottom panels) sessions. Note the more pronounced increase in intracortical facilitation (ICF) after intermittent theta burst stimulation (iTBS) in the HIIT session in young compared to older adults.

### Corticomotor excitability (Figure 2F)

As CME was normalised to the post-exercise time point, we conducted a 2 × 2 (Age x Session) mixed ANOVA, using the average of the three iTBS time points. This analysis revealed a main effect of Session, *F*(1,38 = 9.94, *p* = .003, 17_p_^2^ = 0.21), such that CME was higher following iTBS in the HIIT condition compared to the rest condition (*p* = .003, Cohen’s *d* = 0.72). There was no Session x Age interaction, *F*(1,38 = 0.21, *p* = .649, 17_p_^2^ = 0.006), nor was there a between-subjects effect of Age, *F*(1,38 = 0.05, *p* = .817, 17_p_^2^ = 0.001).

To determine whether exercise or rest alone influenced the amplitude of MEPs to single-pulse TMS, we also conducted a mixed ANOVA using the non-normalised CME data from the baseline and post-exercise time points. Consistent with previous evidence (Smith *et al*., 2014; Andrews *et al*., 2020; Curtin *et al*., 2023), this analysis revealed no main effect of Session, *F*(1,38) = 1.34, *p* = .255, 17_p_^2^ = 0.03, Time, *F*(1,38) = 0.21, *p* = .648, 17_p_^2^ = 0.006, or a Session x Time interaction, *F*(1,38) = 1.04, *p* = .315, 17_p_^2^ = 0.03. There was also no main effect of Age, *F*(1,38) = 0.73, *p* = .397, 17_p_^2^ = 0.02, nor were there any interactions between Session x Age *F*(1,38) = 0.13, *p* = .721, 17_p_^2^ = 0.003, Time x Age, *F*(1,38) = 1.04, *p* = .315, 17_p_^2^ = 0.03, or a Session x Time x Age interaction, *F*(1,38) = 0.01, *p* = .906, 17_p_^2^ < 0.001.

### Negative relationship between age and motor cortex plasticity

To further examine the relationship between ageing and modulation of the E:I balance following exercise and iTBS, we conducted a Pearson correlation and linear regression. This analysis revealed a negative correlation between age and delta E:I balance, *F*(1,18) = 6.56, *R^2^* = .27, *r* = – .52, *p* = .020), such that older age was associated with a reduction in the shift towards reduced excitation and increased inhibition following exercise and iTBS (Figure 4). There was no significant relationship between age and delta E:I balance in our younger group, *F*(1,18) = 0.05, *R^2^* = .003, *r* = – .05, *p* = .832).

**Figure 4.**
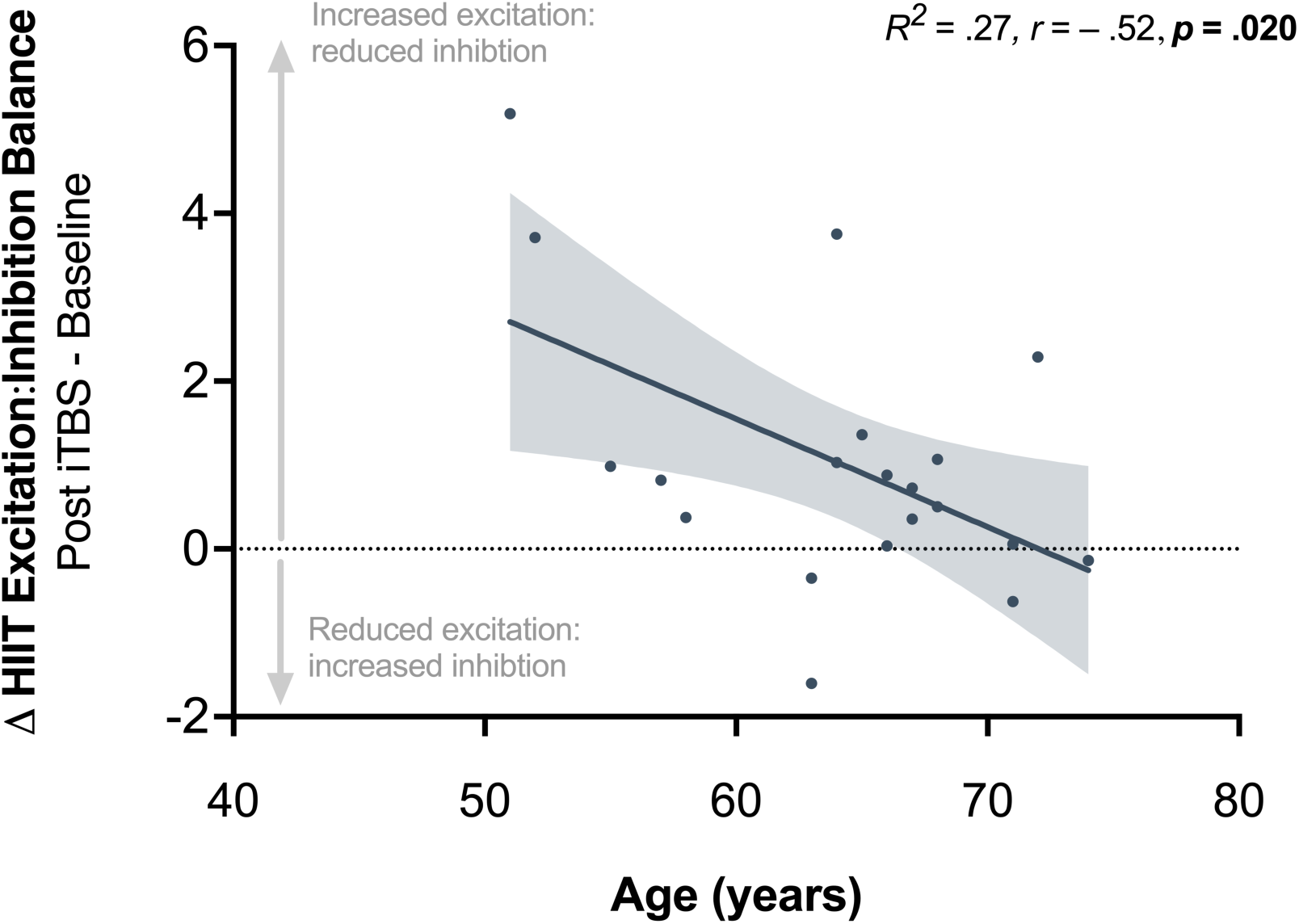
Scatterplot representing the relationship between older age and changes in cortical excitation:inhibition (E:I) following high-intensity exercise (HIIT) and intermittent theta burst stimulation (iTBS). Older age was negatively associated with the change in the excitation:inhibition balance (*R^2^* = .27, *r* = – .52, *p* = .020). Shaded error band denotes the 95% confidence intervals for the line of best fit.

### Background muscle activity

To verify that subthreshold muscle activity (i.e., ≤ 10 μV) did not influence CME, SICI, or ICF across sessions or age groups, separate ANOVAs were run with a between-subject factor of Age (Young, Old) and within-subjects factors of Session (HIIT, rest) and Time (Baseline, Post-Exercise, Post-iTBS 5, Post-iTBS 15, Post-iTBS 25). There was a main effect of Time for CME (*p* = .022), SICI (*p* < .0001), and ICF (*p* = .005), as well as a Session x Time interaction for each TMS variable (all *p ≤* .004), such that root-mean-square EMG decreased following exercise, but this was likely due to reduced electrode impedance as opposed to a meaningful physiological change. Importantly, these analyses revealed no main or interaction effects of Age or Session for our three outcome variables (all *p* ≥ .130), indicating there were no differences in pre-trigger EMG activity across age groups or exercise conditions (Figure 5).

**Figure 5.**
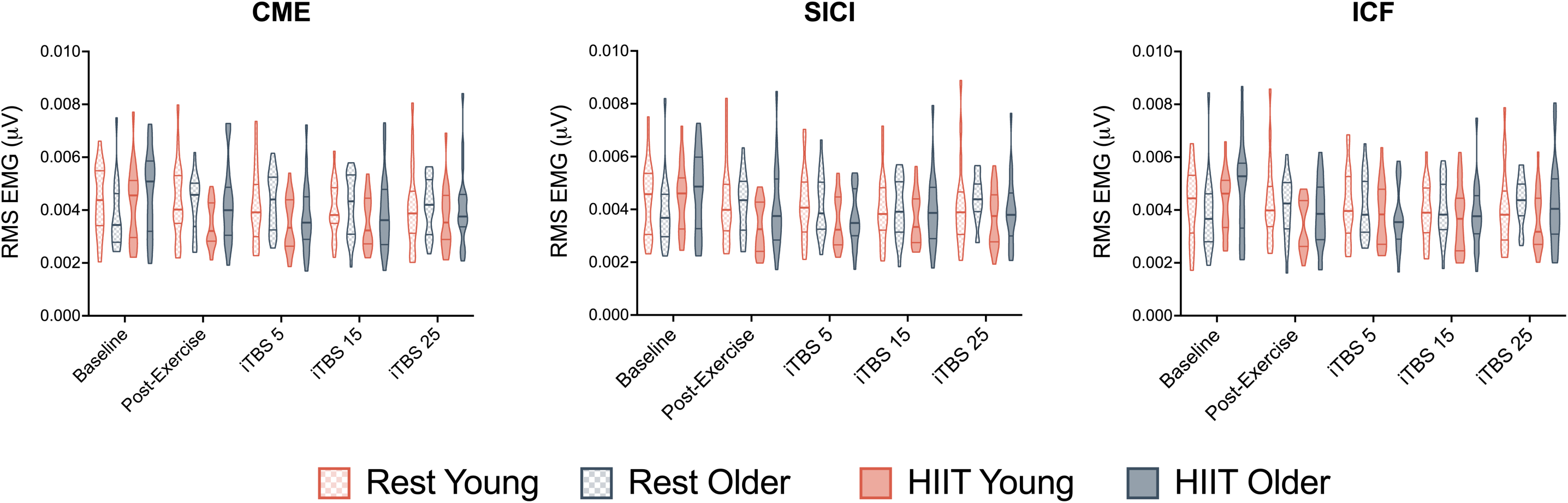
Background muscle activity. There were no differences in pre-trigger EMG activity across age groups or exercise conditions (all *p* ≥ .130). RMS = Root-mean-square, EMG = electromyography, CME = corticomotor excitability, SICI = short-interval intracortical inhibition, ICF = intracortical facilitation, iTBS = intermittent theta burst stimulation, HIIT = high-intensity interval exercise.

## Discussion

We examined the effects of acute cardiorespiratory exercise on motor cortical plasticity in young and older adults. In both groups, exercise shifted the excitation:inhibition (E:I) balance towards an increase in glutamatergic excitation and a reduction in GABAergic inhibition. However, in line with our expectations, older adults showed a reduced plasticity response to intermittent theta burst stimulation (iTBS) following exercise, relative to younger adults. Furthermore, we found that within our older sample, increased age was associated with an attenuated shift in the E:I balance following exercise and iTBS. Taken together, these results demonstrate that the capacity for HIIT exercise to enhance motor cortex plasticity declines with age.

### Ageing attenuated the enhancement of plasticity by exercise

The main finding of our study was that ageing attenuated exercise-induced changes in the E:I balance following iTBS. This result is consistent with the Bienenstock-Cooper-Munro (BCM) theory of bidirectional synaptic plasticity (Bienenstock *et al*., 1982). According to this theory, the capacity for plasticity is modified by previous activity within the same neuron or neuronal network. This modification enables activity to be maintained within a stable physiological range, i.e., to minimise excessive potentiation or depression and is referred to as homeostatic metaplasticity (Abraham, 2008; Müller-Dahlhaus & Ziemann, 2015). If exercise promotes a cortical environment that is conducive to LTP plasticity, then the BCM theory predicts an increased threshold for the subsequent induction of LTP plasticity following iTBS. In support of this account, we found that while exercise promoted changes in motor cortical excitatory and inhibitory networks, these effects were attenuated in older adults following iTBS.

An interesting question raised by such findings is why metaplastic interactions were different in young and older adults. Whereas the after-effects of iTBS were attenuated in older adults following exercise, younger individuals showed further facilitation in plasticity following iTBS. We have previously argued that the results in younger adults are consistent with non-homeostatic plasticity (Müller-Dahlhaus & Ziemann, 2015; Andrews *et al*., 2020). One interpretation of these new findings is that younger adults have a larger range for plasticity, whereas as in older adults, this range may be more constrained and homeostatically regulated (i.e., a lower ‘ceiling’ for plasticity). Although we are not aware of any further evidence examining this question in the context of exercise, previous non-invasive brain stimulation studies have also shown age-related differences in metaplasticity responses (see e.g., Opie *et al*., 2017), suggesting that metaplastic interactions differ in young and older adults. Ageing is associated with declines in many of the neurochemical systems known to be important for regulating plasticity (e.g., dopamine, brain-derived neurotrophic factor, and NMDA receptor) (Backman *et al*., 2010; Erickson *et al*., 2010; Oh *et al*., 2016; Karrer *et al*., 2017; Karalija *et al*., 2022), and it is possible that these changes may contribute to alterations in the capacity for metaplasticity.

An alternative explanation for these results is that the effects of iTBS are less reliable in older, compared to younger, adults. This is a relevant consideration given that previous studies have shown age-related differences in the response to iTBS, as well as other forms of non-invasive brain stimulation (Muller-Dahlhaus *et al*., 2008; Zimerman & Hummel, 2010; Freitas *et al*., 2011). However, we consider this explanation unlikely given that the response to iTBS was not significantly different between young and older groups in the rest condition (see e.g., Figure 2B). This latter finding is consistent with an emerging idea that age-dependent declines in motor cortical plasticity may not be as strong as indicated by early studies (see (Semmler *et al*., 2021) for further discussion).

It is notable that exercise-induced plasticity responses varied even within our older group (see Figure 4). Although we had a relatively small number of participants in the lower age range of the older adult group (i.e., 50 to 60 years), these participants showed plasticity responses more akin to younger adults. People in this age bracket generally had fewer impediments to engaging in high-intensity interval exercise, and likely achieved greater physical exertion, on average, than those over the age of 70 years. The variation in plasticity responses among the older population warrants further investigation using stratified samples with larger group sizes. In particular, it would be interesting to investigate how multiple bouts of exercise or changes in cardiorespiratory fitness affect the plasticity response in older adults. However, overall, these findings are promising as they suggest that exercise-induced changes in plasticity are possible in early and middle ageing.

### Exercise modulates the excitation:inhibition balance in both young and older adults

Consistent with an accumulating body of literature (Singh *et al*., 2014; Smith *et al*., 2014; Mooney *et al*., 2016; Neva *et al*., 2017; Stavrinos & Coxon, 2017; Coxon *et al*., 2018; Morris *et al*., 2020; Curtin *et al*., 2023), we showed that exercise alone (i.e., without iTBS) shifted the E:I balance towards increased excitation and reduced inhibition in younger adults. To date, the effects of healthy ageing on this relationship remain poorly understood. We address this knowledge gap, by showing that a single bout of high-intensity exercise also shifted the E:I balance in the same direction in older adults. A recent study in older adults (Neva *et al*., 2022) did not show significant changes in SICI or ICF following an acute bout of exercise at the group level; however, the pattern of results was consistent with the present study showing a shift in excitatory and inhibitory circuits. The authors argued that their results were obscured by large individual differences associated with their between-subjects design, as well as by the lack of a healthy young comparison group. Critically, we addressed both limitations in the current study using a mixed design, allowing us to make direct age comparisons following exercise and rest conditions.

Evidence from TMS and magnetic resonance spectroscopy (MRS) studies provide insights into the potential behavioural implications of these findings. Such studies have identified changes in motor cortical GABAergic inhibition (Fujiyama *et al*., 2012; Heise *et al*., 2013; Cuypers *et al*., 2020) and glutamatergic excitation (Opie *et al*., 2020) as key mechanisms underpinning motor function in older age. Our results show that it is possible to directly modulate these circuits with exercise, suggesting that exercise may offer a way to mitigate associated age-related decline.

### Limitations

A limitation of the current study was that our older sample were generally high functioning and physically active. Although this sample is representative of a significant portion of the older population, many other older adults are less physically active and have one or more functional dependencies. On the one hand, excluding these participants increases our confidence that our results are not due to any underlying pathologies. On the other, our results cannot be generalised to all members of the ageing population (i.e., those that are more sedentary). For example, based on evidence from young adults (see e.g., Cirillo *et al*., 2009), it is conceivable that older adults with lower cardiorespiratory fitness may exhibit different plasticity responses to those with higher fitness. Further investigation is critical to address the impact of physical fitness, especially given the possibility that these people may have the most to gain from exercise in terms of slowing or mitigating losses in their motor function.

## Conclusion

In both young and older healthy adults, exercise shifted the excitation:inhibition (E:I) balance towards an increase in glutamatergic excitation and a reduction in GABAergic inhibition. In contrast, however, older adults showed an attenuated plasticity response to intermittent theta burst stimulation following exercise, relative to younger adults. Our findings indicate that metaplastic interactions following exercise are altered in ageing and suggest that a preceding bout of high-intensity interval exercise may be less effective for enhancing primary motor cortex plasticity in older adults. Our findings align with the hypothesis that the capacity for cortical plasticity is reduced in older age.

## Abbreviations

E:I: excitation:inhibition
ICF: intracortical facilitation
iTBS: intermittent theta burst stimulation
SICI: short-interval intracortical inhibition
TMS: transcranial magnetic stimulation

## Author contributions

All data was collected at Monash University, Clayton campus, Victoria, Australia. **Dylan Curtin**: Conceptualization, Data acquisition, Formal analysis, Writing - Original Draft, Visualisation. **Claire J. Cadwallader**: Conceptualization, Data acquisition, Formal analysis, Investigation, Writing - Review & Editing. **Eleanor M. Taylor**: Conceptualization, Data acquisition, Writing - Review & Editing. **Sophie C Andrews**: Conceptualization, Writing - Review & Editing, Funding acquisition, **Julie C Stout:** Conceptualization, Writing - Review & Editing, Funding acquisition, **Joshua J. Hendrikse**: Conceptualization, Data acquisition, Writing - Review & Editing, **Trevor T-J. Chong**: Conceptualization, Writing - Review & Editing, Funding acquisition. **James P. Coxon**: Conceptualization, Data acquisition, Formal analysis, Writing - Review & Editing, Visualisation, Funding acquisition.

## Acknowledgements

We thank Tamar de Moel for her support with data collection, and Emily Brooks, Sarah Wallis, and Sarah Cohen for their assistance with participant recruitment.

## References

Abraham WC. (2008). Metaplasticity: tuning synapses and networks for plasticity. Nat Rev Neurosci 9, 387.

Andrews SC, Curtin D, Coxon JP & Stout JC. (2022). Motor cortex plasticity response to acute cardiorespiratory exercise and intermittent theta-burst stimulation is attenuated in premanifest and early Huntington’s disease. Sci Rep 12, 1104.

Andrews SC, Curtin D, Hawi Z, Wongtrakun J, Stout JC & Coxon JP. (2020). Intensity matters: High-intensity interval exercise enhances motor cortex plasticity more than moderate exercise. Cereb Cortex.

Backman L, Lindenberger U, Li SC & Nyberg L. (2010). Linking cognitive aging to alterations in dopamine neurotransmitter functioning: recent data and future avenues. Neurosci Biobehav Rev 34, 670–677.

Bhandari A, Radhu N, Farzan F, Mulsant BH, Rajji TK, Daskalakis ZJ & Blumberger DM. (2016). A meta-analysis of the effects of aging on motor cortex neurophysiology assessed by transcranial magnetic stimulation. Clin Neurophysiol 127, 2834–2845.

Bienenstock EL, Cooper LN & Munro PW. (1982). Theory for the development of neuron selectivity: orientation specificity and binocular interaction in visual cortex. J Neurosci 2, 32–48.

Borg G. (1970). Perceived exertion as an indicator of somatic stress. Scand J Rehabil Med 2, 92–98.

Cadwallader CJ, Steiniger J, Cooper PS, Zhou SH, Hendrikse J, Sumner RL, Kirk IJ, Chong TT & Coxon JP. (2023). Acute exercise as a modifier of neocortical plasticity and aperiodic activity in the visual cortex. Sci Rep 13, 7491.

Cirillo J, Lavender AP, Ridding MC & Semmler JG. (2009). Motor cortex plasticity induced by paired associative stimulation is enhanced in physically active individuals. J Physiol 587, 5831–5842.

Coxon JP, Cash RFH, Hendrikse JJ, Rogasch NC, Stavrinos E, Suo C & Yucel M. (2018). GABA concentration in sensorimotor cortex following high-intensity exercise and relationship to lactate levels. J Physiol 596, 691–702.

Coxon JP, Peat NM & Byblow WD. (2014). Primary motor cortex disinhibition during motor skill learning. J Neurophysiol 112, 156–164.

Craig CL, Marshall AL, Sjöström M, Bauman AE, Booth ML, Ainsworth BE, Pratt M, Ekelund U, Yngve A, Sallis JF & Oja P. (2003). International physical activity questionnaire: 12-Country reliability and validity. Med Sci Sports Exerc 35, 1381–1395.

Curtin D, Taylor EM, Bellgrove MA, Chong TT & Coxon JP. (2023). D2 receptor blockade eliminates exercise-induced changes in cortical inhibition and excitation. Brain Stimul 16, 727–733.

Cuypers K, Verstraelen S, Maes C, Hermans L, Hehl M, Heise KF, Chalavi S, Mikkelsen M, Edden R, Levin O, Sunaert S, Meesen R, Mantini D & Swinnen SP. (2020). Task-related measures of short-interval intracortical inhibition and GABA levels in healthy young and older adults: A multimodal TMS-MRS study. Neuroimage 208, 116470.

Di Lorito C, Long A, Byrne A, Harwood RH, Gladman JRF, Schneider S, Logan P, Bosco A & van der Wardt V. (2021). Exercise interventions for older adults: A systematic review of meta-analyses. J Sport Health Sci 10, 29–47.

Edvardsen E, Hem E & Anderssen SA. (2014). End criteria for reaching maximal oxygen uptake must be strict and adjusted to sex and age: a cross-sectional study. PLoS One 9, e85276.

Erickson KI, Donofry SD, Sewell KR, Brown BM & Stillman CM. (2022). Cognitive Aging and the Promise of Physical Activity. Annu Rev Clin Psychol 18, 417–442.

Erickson KI, Prakash RS, Voss MW, Chaddock L, Heo S, McLaren M, Pence BD, Martin SA, Vieira VJ, Woods JA, McAuley E & Kramer AF. (2010). Brain-derived neurotrophic factor is associated with age-related decline in hippocampal volume. J Neurosci 30, 5368–5375.

Erickson KI, Voss MW, Prakash RS, Basak C, Szabo A, Chaddock L, Kim JS, Heo S, Alves H, White SM, Wojcicki TR, Mailey E, Vieira VJ, Martin SA, Pence BD, Woods JA, McAuley E & Kramer AF. (2011). Exercise training increases size of hippocampus and improves memory. Proc Natl Acad Sci U S A 108, 3017–3022.

Freitas C, Perez J, Knobel M, Tormos JM, Oberman L, Eldaief M, Bashir S, Vernet M, Peña-Gómez C & Pascual-Leone A. (2011). Changes in cortical plasticity across the lifespan. Front Aging Neurosci 3, 1–8.

Fujiyama H, Hinder MR, Schmidt MW, Tandonnet C, Garry MI & Summers JJ. (2012). Age-related Differences in Corticomotor Excitability and Inhibitory Processes during a Visuomotor RT Task. J Cogn Neurosci 24, 1253–1263.

Heise KF, Zimerman M, Hoppe J, Gerloff C, Wegscheider K & Hummel FC. (2013). The aging motor system as a model for plastic changes of GABA-mediated intracortical inhibition and their behavioral relevance. J Neurosci 33, 9039–9049.

Hendrikse J, Thompson S, Suo C, Yücel M, Rogasch NC & Coxon JP. (2021). The effects of multi-day rTMS and cardiorespiratory fitness on working memory and local GABA concentration. Neuroimage: Reports 1.

Hermans L, Levin O, Maes C, van Ruitenbeek P, Heise KF, Edden RAE, Puts NAJ, Peeters R, King BR, Meesen RLJ, Leunissen I, Swinnen SP & Cuypers K. (2018). GABA levels and measures of intracortical and interhemispheric excitability in healthy young and older adults: an MRS-TMS study. Neurobiol Aging 65, 168–177.

Huang Y-Z, Rothwell JC, Edwards MJ & Chen R-S. (2007). Effect of physiological activity on an NMDA-dependent form of cortical plasticity in human. Cereb Cortex 18, 563–570.

Huang YZ, Edwards MJ, Rounis E, Bhatia KP & Rothwell JC. (2005). Theta burst stimulation of the human motor cortex. Neuron 45, 201–206.

Ji L, Steffens DC & Wang L. (2021). Effects of physical exercise on the aging brain across imaging modalities: A meta-analysis of neuroimaging studies in randomized controlled trials. Int J Geriatr Psychiatry 36, 1148–1157.

Karalija N, Johansson J, Papenberg G, Wahlin A, Salami A, Kohncke Y, Brandmaier AM, Andersson M, Axelsson J, Riklund K, Lovden M, Lindenberger U, Backman L & Nyberg L. (2022). Longitudinal Dopamine D2 Receptor Changes and Cerebrovascular Health in Aging. Neurology 99, e1278–e1289.

Karrer TM, Josef AK, Mata R, Morris ED & Samanez-Larkin GR. (2017). Reduced dopamine receptors and transporters but not synthesis capacity in normal aging adults: a meta-analysis. Neurobiol Aging 57, 36–46.

Kolasinski J, Hinson EL, Divanbeighi Zand AP, Rizov A, Emir UE & Stagg CJ. (2019). The dynamics of cortical GABA in human motor learning. J Physiol 597, 271–282.

Levin O, Fujiyama H, Boisgontier MP, Swinnen SP & Summers JJ. (2014). Aging and motor inhibition: a converging perspective provided by brain stimulation and imaging approaches. Neurosci Biobehav Rev 43, 100–117.

Maass A, Duzel S, Goerke M, Becke A, Sobieray U, Neumann K, Lovden M, Lindenberger U, Backman L, Braun-Dullaeus R, Ahrens D, Heinze HJ, Muller NG & Duzel E. (2015). Vascular hippocampal plasticity after aerobic exercise in older adults. Mol Psychiatry 20, 585–593.

Mang CS, Snow NJ, Campbell KL, Ross CJ & Boyd LA. (2014). A single bout of high-intensity aerobic exercise facilitates response to paired associative stimulation and promotes sequence-specific implicit motor learning. J Appl Physiol (1985) 117, 1325-1336.

Marneweck M, Loftus A & Hammond G. (2011). Short-interval intracortical inhibition and manual dexterity in healthy aging. Neurosci Res 70, 408–414.

McGinley M, Hoffman RL, Russ DW, Thomas JS & Clark BC. (2010). Older adults exhibit more intracortical inhibition and less intracortical facilitation than young adults. Exp Gerontol 45, 671–678.

Mooney RA, Cirillo J & Byblow WD. (2017). GABA and primary motor cortex inhibition in young and older adults: a multimodal reliability study. J Neurophysiol 118, 425–433.

Mooney RA, Coxon JP, Cirillo J, Glenny H, Gant N & Byblow WD. (2016). Acute aerobic exercise modulates primary motor cortex inhibition. Exp Brain Res 234, 3669–3676.

Morris TP, Fried PJ, Macone J, Stillman A, Gomes-Osman J, Costa-Miserachs D, Tormos Munoz JM, Santarnecchi E & Pascual-Leone A. (2020). Light aerobic exercise modulates executive function and cortical excitability. Eur J Neurosci 51, 1723–1734.

Muellbacher W, Ziemann U, Wissel J, Dang N, Kofler M, Facchini S, Boroojerdi B, Poewe W & Hallett M. (2002). Early consolidation in human primary motor cortex. Nature 415, 640–644.

Müller-Dahlhaus F & Ziemann U. (2015). Metaplasticity in human cortex. The Neuroscientist 21, 185–202.

Muller-Dahlhaus JF, Orekhov Y, Liu Y & Ziemann U. (2008). Interindividual variability and age-dependency of motor cortical plasticity induced by paired associative stimulation. Exp Brain Res 187, 467–475.

Nasreddine ZS, Phillips NA, Bédirian V, Charbonneau S, Whitehead V, Collin I, Cummings JL & Chertkow H. (2005). The Montreal Cognitive Assessment, MoCA: a brief screening tool for mild cognitive impairment. J Am Geriatr Soc 53, 695–699.

Nes BM, Janszky I, Wisloff U, Stoylen A & Karlsen T. (2013). Age-predicted maximal heart rate in healthy subjects: The HUNT fitness study. Scand J Med Sci Sports 23, 697–704.

Neva JL, Brown KE, Mang CS, Francisco BA & Boyd LA. (2017). An acute bout of exercise modulates both intracortical and interhemispheric excitability. Eur J Neurosci 45, 1343–1355.

Neva JL, Greeley B, Chau B, Ferris JK, Jones CB, Denyer R, Hayward KS, Campbell KL & Boyd LA. (2022). Acute High-Intensity Interval Exercise Modulates Corticospinal Excitability in Older Adults. Med Sci Sports Exerc 54, 673–682.

Oh H, Lewis DA & Sibille E. (2016). The Role of BDNF in Age-Dependent Changes of Excitatory and Inhibitory Synaptic Markers in the Human Prefrontal Cortex. Neuropsychopharmacology 41, 3080–3091.

Oldfield RC. (1971). The assessment and analysis of handedness: The Edinburgh inventory. Neuropsychologia 9, 97–113.

Opie GM, Hand BJ & Semmler JG. (2020). Age-related changes in late synaptic inputs to corticospinal neurons and their functional significance: A paired-pulse TMS study. Brain Stimul 13, 239–246.

Opie GM, Post AK, Ridding MC, Ziemann U & Semmler JG. (2017). Modulating motor cortical neuroplasticity with priming paired associative stimulation in young and old adults. Clin Neurophysiol 128, 763–769.

Peinemann A, Lehner C, Conrad B & Siebner HR. (2001). Age-related decrease in paired-pulse intracortical inhibition in the human primary motor cortex. Neurosci Lett 313, 33–36.

Peters AJ, Liu H & Komiyama T. (2017). Learning in the Rodent Motor Cortex. Annu Rev Neurosci 40, 77–97.

Ridding MC & Ziemann U. (2010). Determinants of the induction of cortical plasticity by non-invasive brain stimulation in healthy subjects. J Physiol 588, 2291–2304.

Rioult-Pedotti M-S, Friedman D, Hess G & Donoghue JP. (1998). Strengthening of horizontal cortical connections following skill learning. Nat Neurosci 1, 230–234.

Rossi S, Hallett M, Rossini PM, Pascual-Leone A, Avanzini G, Bestmann S, Berardelli A, Brewer C, Canli T, Cantello R, Chen R, Classen J, Demitrack M, Di Lazzaro V, Epstein CM, George MS, Fregni F, Ilmoniemi R, Jalinous R, Karp B, Lefaucheur JP, Lisanby S, Meunier S, Miniussi C, Miranda P, Padberg F, Paulus W, Peterchev A, Porteri C, Provost M, Quartarone A, Rotenberg A, Rothwell J, Ruohonen J, Siebner H, Thut G, Valls-Solè J, Walsh V, Ugawa Y, Zangen A & Ziemann U. (2009). Safety, ethical considerations, and application guidelines for the use of transcranial magnetic stimulation in clinical practice and research. Clin Neurophysiol 120, 2008–2039.

Sanes JN & Donoghue JP. (2000). Plasticity and primary motor cortex. In Annu Rev Neurosci, pp. 393-415.

Seidler RD, Bernard JA, Burutolu TB, Fling BW, Gordon MT, Gwin JT, Kwak Y & Lipps DB. (2010). Motor control and aging: links to age-related brain structural, functional, and biochemical effects. Neurosci Biobehav Rev 34, 721–733.

Semmler JG, Hand BJ, Sasaki R, Merkin A & Opie GM. (2021). Age-related changes in motor cortex plasticity assessed with non-invasive brain stimulation: an update and new perspectives. Exp Brain Res 239, 2661–2678.

Singh AM, Duncan RE, Neva JL & Staines WR. (2014). Aerobic exercise modulates intracortical inhibition and facilitation in a nonexercised upper limb muscle. BMC Sports Science, Medicine and Rehabilitation 6.

Smith AE, Goldsworthy MR, Garside T, Wood FM & Ridding MC. (2014). The influence of a single bout of aerobic exercise on short-interval intracortical excitability. Exp Brain Res 232, 1875–1882.

Sports Medicine Australia GoA, Canberra, ACT. (2011). Sports Medicine Australia Pre-exercise Screening System.

Stavrinos EL & Coxon JP. (2017). High-intensity interval exercise promotes motor cortex disinhibition and early motor skill consolidation. J Cogn Neurosci 29, 593–604.

Suppa A, Huang YZ, Funke K, Ridding MC, Cheeran B, Di Lazzaro V, Ziemann U & Rothwell JC. (2016). Ten Years of Theta Burst Stimulation in Humans: Established Knowledge, Unknowns and Prospects. Brain Stimul 9, 323–335.

Tabachnick BG & Fidell LS. (2013). Using Multivariate Statistics. Textbook.

Tanaka H, Monahan KD & Seals DR. (2001). Age-predicted maximal heart rate revisited. J Am Coll Cardiol 37, 153–156.

Valenzuela PL, Saco-Ledo G, Morales JS, Gallardo-Gomez D, Morales-Palomo F, Lopez-Ortiz S, Rivas-Baeza B, Castillo-Garcia A, Jimenez-Pavon D, Santos-Lozano A, Del Pozo Cruz B & Lucia A. (2023). Effects of physical exercise on physical function in older adults in residential care: a systematic review and network meta-analysis of randomised controlled trials. Lancet Healthy Longev.

Zimerman M & Hummel FC. (2010). Non-invasive brain stimulation: enhancing motor and cognitive functions in healthy old subjects. Front Aging Neurosci 2, 149.

